# Identifying Charismatic Bird Species and Traits with Community Science Observations

**DOI:** 10.1101/2021.06.05.446577

**Authors:** Sara Stoudt, Benjamin R. Goldstein, Perry De Valpine

## Abstract

Identifying which species are perceived as charismatic can improve the impact and efficiency of conservation outreach, as charismatic species receive more conservation funding and have their conservation needs prioritized (9; 17; 13). Sociological experiments studying animal charisma have relied on stated preferences to find correlations between hypothetical “willingness to pay” or “empathy” for a species’ conservation and species’ size, color, and aesthetic appeal (51; 13; 16). Recognizing the increasing availability of digital records of public engagement with animals that reveal preferences, an emerging field of “culturomics” uses Google search results, Wikipedia article activities, and other digital modes of engagement to identify charismatic species and traits (46; 31; 10; 41). In this study, we take advantage of community science efforts as another form of digital data that can reveal observer preferences. We apply a multi-stage analysis to ask whether opportunistic birders contributing to iNaturalist engage more with larger, more colorful, and rarer birds relative to a baseline, from eBird contributors, approximating unbiased detection. We find that body mass, color contrast, and range size all predict over-representation in the opportunistic dataset. We also find evidence that, across 473 modeled species, 52 species are significantly overreported and 158 are significantly underreported, indicating a wide variety of species-specific effects. Understanding which birds are charismatic can aid conservationists in creating impactful outreach materials and engaging new naturalists. The quantified differences between two prominent community science efforts may also be of use for researchers leveraging the data from one or both of them to answer scientific questions of interest.

## 1. Introduction

Birds have received special attention in conservation (21; 48), and investigations into stated preferences for birds found that species traits—color, pattern, and shape—influence their perceived charisma (9; 37; 38). Others have taken advantage of revealed-preference data for birds from the volume of Google search results and the Common Breeding Bird Monitoring Scheme (62) and eBird data (50) to similarly investigate the public’s perception of different birds.

Online community science platforms, which collect data contributed by volunteers, provide a more direct way to study public perception of species in the wild. Community scientists, sometimes called “citizen scientists,” volunteer contributions to scientific databases as self-guided, non-professionals. Two biodiversity platforms in particular are of great interest for investigating bird charisma on a large scale. eBird, an app for hobbyist birders that has generated one of the world’s largest biodiversity databases, has recorded over 550 million records in North America to date (22). Many of these records come from over 46 million “complete checklists” (20) and thus represent a rigorous reporting protocol, with reliable information on when species were not seen alongside when they were seen as well as the inclusion of sampling metadata (18). Another popular platform is iNaturalist, a nature app designed to encourage public engagement with all species. The primary goal of iNaturalist is to “connect people to nature” (28). The app allows any observation of any species at any time or place to be entered, so reporting rates depend on relative interest in different species.

Biodiversity records such as those aggregated on online community science platforms are important for informing species distribution models (54; 14; 43; 56). Complete checklists from eBird are lauded as appropriate for species distribution modeling (11; 53), whereas opportunistic records such as from iNaturalist are known to contain particular biases (29; 40). These biases are often characterized as noise, but they may actually contain a strong signal of the habits and preferences of opportunistic naturalists. In this paper, we estimate those biases in relation to eBird and thereby analyze public interest in species traits such as size, color contrast, phylogenetic relatedness, and taxonomic order. By harnessing two huge but very different community science datasets, we gain insight into human interest in biodiversity as encountered in the wild.

We construct a conceptual model to relate eBird and iNaturalist’s data-generating processes and show how they can be studied to characterize observer biases and preferences (Figure 1). Imagine an iNaturalist user who sees a bird, takes a picture of it, and submits the photo to iNaturalist. For this event to occur, three separate conditions must be satisfied. First, the species must be present in the environment. We call this condition the “presence” filter, and characterizing this process is the main goal of most species distribution models that use community science data. Second, the observer must see the species–this is the “detectability” filter, which is controlled for in ecological studies as imperfect detection. Were this eBird, the process would stop here, because all detected birds must be reported under the complete-checklist protocol. However, in iNaturalist, a third event must occur: having seen the species, the observer must then make the choice to record an audio clip or, far more commonly, take a photo and upload it to iNaturalist. We call this the “interest” filter. Characterizing the interest filter, visible as deviations in iNaturalist reporting rates relative to eBird, is the main goal of this study.

**Figure 1.**
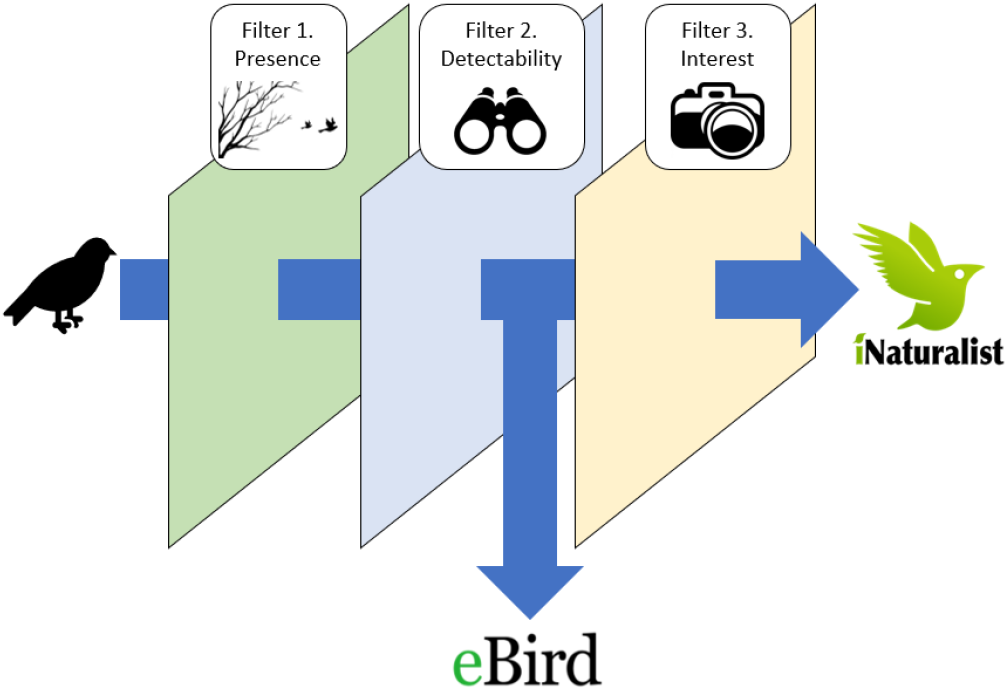
Community science reporting as a filtering process. eBird data passes through two conceptual filters: (1) presence and (2) detectability. In iNaturalist, a hypothetical report must pass through an additional filter, (3) interest.

## 2. Materials and methods

### 2.1. eBird and iNaturalist pre-processing and data structures

In August 2020 we downloaded the eBird Basic Dataset and extracted all complete checklists. We also obtained all iNaturalist research grade observations of bird species from the Global Biodiversity Information Facility (23). For both datasets, we considered only observations made in the contiguous 48 United States and Washington, D.C. on or before December 31, 2019. We excluded checklists obtained in 2020 or later to avoid accidentally capturing changes in observer behavior related to the COVID-19 pandemic (25; 15). We associated species across the two datasets using the R package taxalight (8).

We aggregated observations to a spatial grid of regular hexagonal cells covering the contiguous U.S. such that spatial grid cells had a long radius (center to vertex) of 20 kilometers. In each “hex,” a count was obtained for the number of times each species was detected. The total number of observations in each hex was calculated as the sum of the species observation events in that hex. For consistency with iNaturalist, an eBird checklist that reported more than one individual of a species was counted as a series of separate species observation events (a checklist with three species reported constituted three observations). By aggregating over time we assumed that the primary variation is spatial. We accounted for secondary variation elsewhere in the modeling approach.

We sub-selected species according to the following criteria. First, to capture spatial variability in sampling, we only considered species that were observed one or more times in at least 100 different hexes. Second, we only considered species in the EltonTraits database (60; 58). We eliminated “pelagic specialists” according to EltonTraits, expecting the sampling process generating these data to be fundamentally different from that of terrestrial birds. 482 North American bird species met these criteria.

### 2.2. Species-level spatial analysis

We first estimated a typical overreporting index characterizing each species. The unit of analysis for the first stage was *y_ijk_*, the count of reports of species *j* reported at the *i*th spatial hex in dataset *k* (either eBird or iNaturalist). We modeled the number of “successes” in a binomial random draw *y_ijk_* as *y_ijk_* ~ *Binom*(*C_ik_, R_ijk_*). Here *C_ik_* is the known number of “attempts,” which we define for the presence-only iNaturalist data to be the total number of iNaturalist observations across all bird species in spatial hex *i* and for the eBird checklist data to be the total number of species observation events on all of the eBird checklists in spatial hex *i*. (Note that for eBird, a single checklist on which 5 different species are observed would be considered five species observation events, rather than one, in order to match the iNaturalist sampling schema.) *R_ijk_* is the reporting rate which we model as a function of location *i* for each species *j* and dataset *k* combination.

We used a quasi-binomial generalized additive model (GAM) with a logit link to capture spatial variability via a multidimensional tensor-product smooth of the longitude and latitude coordinates of the hexes (61). The motivation for this approach was twofold. First, we anticipated that many differences between the datasets could be due to spatial heterogeneity in sampling. Both eBird and iNaturalist are highly spatially variable with their own hotspots—for example, eBird is most densely used near its base in Cornell, NY, and on the East Coast, while iNaturalist has high density regions in California and Texas (19; 27). For this reason alone we expect that reporting rates would vary by dataset, and so a spatially explicit analysis is called for. A spatially explicit approach is also necessary since user habits may themselves be spatially non-independent. To obtain accurate confidence intervals on parameter estimates, spatial autocorrelation in the data-generating process must be accounted for. Second, we anticipated extra-binomial, non-spatial variability across units (e.g. temporal, weather conditions, observer variability, etc.). We chose a quasi-binomial approach as a way to account for this in the uncertainty quantification.

A GAM was fit for each species and dataset combination. The basis dimensions were chosen to be 20 knots by 20 knots. We fit all models and then iteratively increased each dimension by 5 until the model passed a hypothesis test of whether the basis dimension for a smooth was adequate using a p-value cutoff *α* = 0.1 (given in the R function mgcv::gam.check).

From the quasi-binomial GAMs, we obtained estimates of the spatially smooth surface of the reporting rate at each hex, in each dataset. We calculated the overreporting index as the median predicted difference in log-scale reporting rates across hexes for each species. To obtain accurate confidence intervals on the overreporting index, we used a parametric bootstrap approach, making random draws of the spatial surface and recomputing the index each time, to obtain an estimate of uncertainty for each index. Each species’ overreporting index represents the typical deviation in the iNaturalist reporting rate relative to the eBird baseline for that species.

To assess which overreporting indices were significantly different from zero, we used a p-value threshold that was adjusted to account for the fact that we made multiple comparisons (one for each species). We used a false discovery rate controlling method to ensure that across comparisons the false positive rate was no more than 0.05 (7).

### 2.3. Cross-species meta-analysis

To answer our main question, whether species traits can help explain these differences in birder engagement, we used a meta-analysis to explain patterns in the overreporting indices. A meta-analysis allowed us to propagate the uncertainty estimated in the first stage of the analysis (57). The median differences in reporting rates for all of the species, along with their standard errors, became the response in this stage of the analysis.

We hypothesized two types of charisma-associated bias that could drive differences between the datasets. The first, photogenic bias, refers to sampling differences due to aesthetic preferences of observers. One component of photogenic bias relates to relevant aspects of species charisma, such as size and color. To investigate the effect of size, we retrieved species’ log mass from the EltonTraits dataset (58). To represent how colorful or striking a bird is, we used an index of maximum color contrast originally developed by Schuetz and Johnston (50). Another component of photogenic bias is logistical: particular bird species might be overreported by virtue of being easier to document rather than more charismatic. Since iNaturalist observations are almost always associated with photos this could lead to a difference between the datasets. The second type of bias, novelty bias, may occur where users in iNaturalist preferentially report species that are new to them or that they see infrequently. We used two covariates as proxies of different aspects of rarity: the number of hexes a species is reported in (a proxy for size of effective range) and the proportion of all eBird checklists where the species was found (a spatially agnostic proxy for overall prevalence). We centered and scaled these covariates for log mass, maximum color contrast, log range size, and log prevalence.

In the meta-analysis we also incorporated phylogenetic structure to account for the possibility that phylogenetically closer species have more similar reporting indices due to evolutionary non-independence of unmodeled but important traits (1; 36). We obtained multiple phylogenetic trees from BirdTree.org (30). We then obtained a consensus tree including branch edges using the R package phytools (47). Finally we computed a variance-covariance matrix based on this consensus tree using the R package ape (45). We allowed for both a random effect for species with this variance-covariance structure and an unstructured random effect for species, similar to an approach of creating a mixture of a phylogenetic covariance matrix and the identity matrix (44).

We fit a second meta-analysis including only the effect of taxonomic order and excluding phylogenetic structure to obtain estimates of each order’s mean overreporting index with properly propagated error (4; 33).

Models for nine species failed to fit (see Appendix) and were therefore dropped from the second stage analysis. Three of these failed to pass a test for adequate knots with a basis dimension of 35 by 35, above which computation became infeasible. Six of these failed to converge in under 24 hours, which was chosen as a practical cutoff.

We removed 49 species that had overreporting indices outside the range −10 to 10 from the meta-analysis stage. Values less than −10 or greater than 10 arose in cases where, among the union of hexes where a species was reported in either dataset, one dataset reported no observations in over half of those hexes. Because the reporting index uses median differences in log reporting rates, it could not be reliably estimated, nor its uncertainty reliably quantified for the meta-analysis step, in these cases. After these two filters, 424 species were included. To test the sensitivity of results, we replicated the meta-analysis including the extreme overreporting indices.

## 3. Results

We predicted an overreporting index with uncertainty for each of 473 species of interest. Even when controlling the false discovery rate, 210 species had overreporting indices significantly different from zero, giving evidence that iNaturalist and eBird reporting rates were meaningfully different for many species. A significantly negative overreporting index means iNaturalist observers were uninterested, while a significantly positive one means they were interested. Figure 2 shows the most extreme over- and under-reported birds. The most overreported birds are each some combination of large (wild turkey), well-known for their appeal (burrowing owl) or considered especially beautiful (Indian peafowl). The least overreported birds follow less of an obvious pattern, though they tend to be smaller or associated with near-ocean foraging.

**Figure 2.**
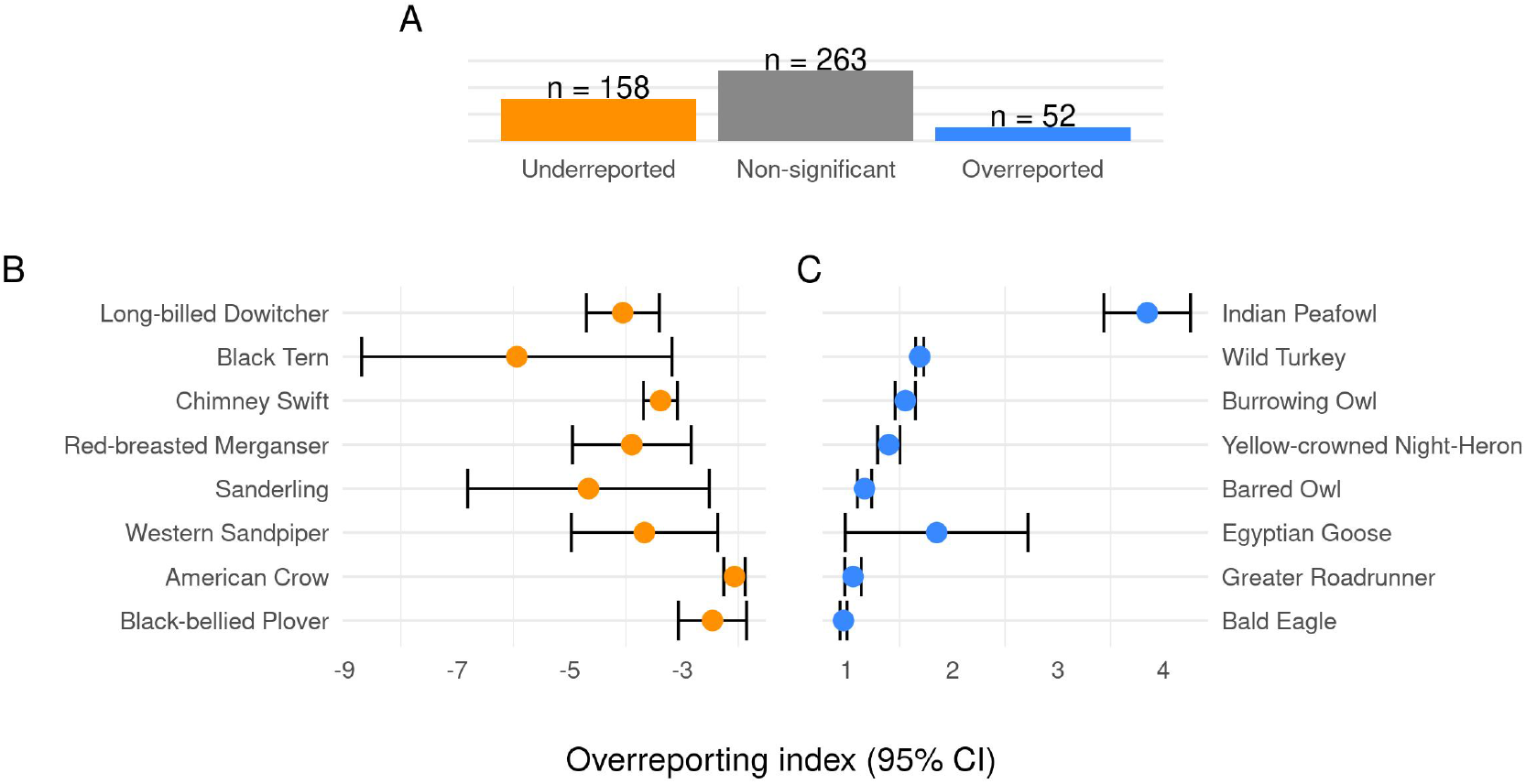
Species overreporting indices. (A) Counts of bird overreporting indices by significance level, controlling the FDR to account for multiple comparisons. The most under-reported species (B) appeared in iNaturalist between 0.002-0.13 as frequently as in eBird, while the most over-reported species (C) appeared in iNaturalist roughly 1.6-45 times as frequently as in eBird. Over-reporting index is median difference in log reporting rate. For each species, the estimate with 95% CI is shown.

We hypothesized that these differences at the species level, represented by the third filter layer (Figure 1), could be driven by a variety of mechanisms. We next investigated whether photogenic bias and novelty bias were present. Figure 3 shows the relationship between overreporting indices and these species traits that relate to the proposed biases.

**Figure 3.**
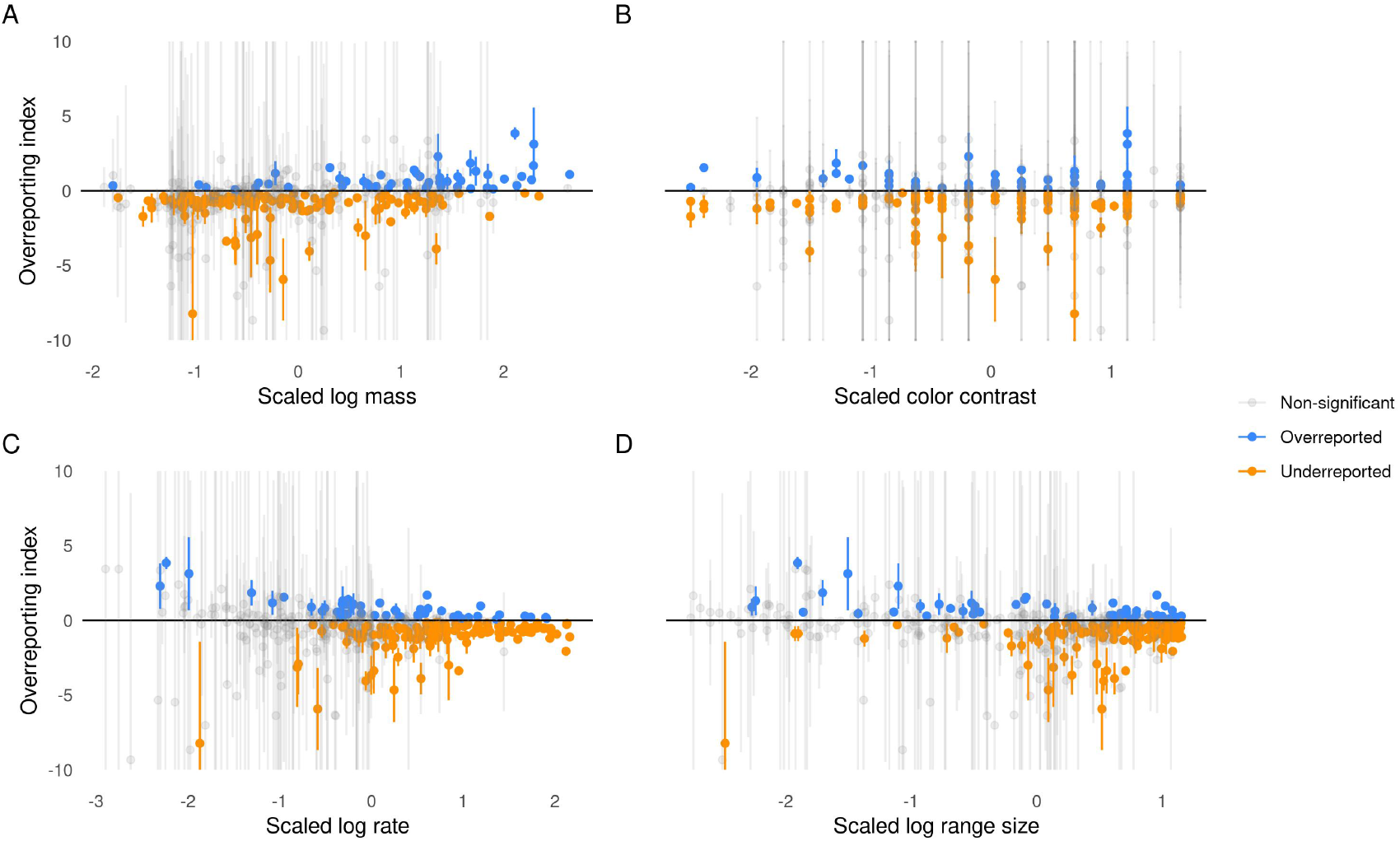
Species overreporting indices plotted against four traits—(A) log mass, (B) color contrast, (C) log prevalance, and (D) log range size—whose associations were studied in a meta-analysis. Vertical error bars show 95% confidence intervals of indices, which do not correspond exactly to the significance test adjusted for multiple testing.

We found a statistically significant relationship between the overreporting index and size, color, and number of hexes containing observations of the species (a proxy for range size) in a second stage meta-analysis. This means that opportunistic birders in the contiguous U.S. chose larger, more colorful, and more range-limited birds more often than would be expected based on the corresponding species’ detection rates in eBird. We found a 0.31 effect size on scaled log mass with a 95% confidence interval of (0.15, 0.47), 0.12 for scaled color (0.04, 0.19), −0.13 for scaled log number of hexes where reported (−0.25, −0.03), and −0.08 for scaled log proportion of eBird checklists where reported (−0.19, 0.03). See Table 1 in the Appendix for more interpretation. The independent random effect dominates the phylogenetically structured random effect in magnitude, suggesting there is not much phylogenetic structure in the residual variance.

**Table 1.** Meta-analysis coefficients and interpretations for species traits once low information outliers were excluded (overreporting indices greater than 10 or less than −10). All covariates except for the proxy of overall prevalence were found to be statistically significant.

| Covariate | Effect Size | Interpretation |
| --- | --- | --- |
| Scaled log mass | 0.31 (0.15, 0.47) | An increase by one standard deviation of bird size on the log scale is associated with an expected 36% increase in the odds of overreporting. |
| Scaled color | 0.12 (0.04, 0.19) | An increase by one standard deviation of bird color contrast is associated with an expected 12% increase in the odds of overreporting. |
| Scaled log number of hexes where reported | -0.13 (-0.25, -0.03) | An increase by one standard deviation of number of hexes on the log scale is associated with an expected 11% decrease in the odds of overreporting. |
| Scaled log proportion of eBird checklists where found | -0.08 (-0.19, 0.03) | An increase by one standard deviation of proportion of eBird checklists where a species is found, on the log scale, is associated with a 8% decrease in the odds of overreporting. |

We tested sensitivity of results to the choice to exclude low information indices with extreme values. Including these indices made the effect of overall prevalence significantly positive, removed the significance of color, and increased the coefficients for mass and the overall prevalence (See Appendix for details). Thus, results are sensitive to an attempt to include unreliable indices for species with limited data. See the Appendix for further discussion of these points.

We accounted for phylogeny at the species level in the meta-analysis, but it can also be helpful to visualize relationships at the order level for improved interpretation. We identified five taxonomic orders whose typical overreporting indices were significantly different from zero after correcting for multiple testing (Figure 4). Owls and gamefowl tend to be relatively large, which may explain their overreporting. However, many waders and gulls are large, and this order is underreported, along with songbirds and doves.

**Figure 4.**
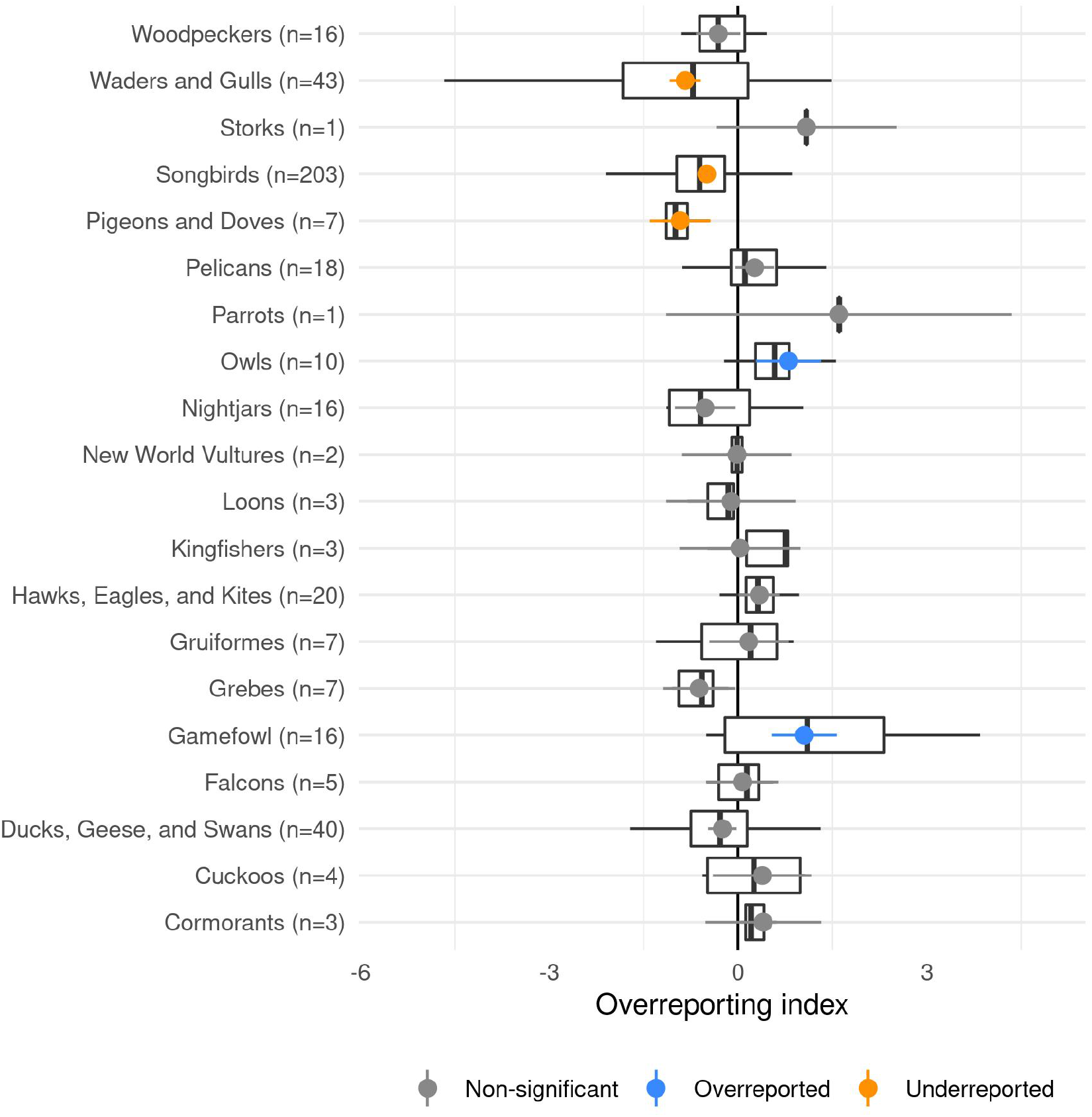
The predicted effect of order, representing the typical overreporting index for that group of species, excluding low-information outliers. Colors indicate the results of a false discovery rate-controlling significance test. Box plots show the range of estimated overreporting indices in each group.

## 4. Discussion

When using community science data to study biodiversity, the inescapable fact of variation between users is usually treated as noise that at best adds uncertainty and at worst causes biased estimation. However, when studying people’s relationships to animals rather than the animals themselves, with an eye towards using this knowledge to inform conservation efforts (35), the paradigm flips, and variation between users becomes the data-generating process of interest. For this task, community science is not merely a source of noisy data, but a unique and invaluable source of information about how members of the public choose to engage with the natural world.

The iNaturalist dataset in particular is ideal for characterizing naturalists’ preferences. Because iNaturalist is open to all living organisms, anything like eBird’s complete checklist protocol is impossible, as an observer will likely see hundreds of plants and dozens of animals for every individual they choose to report. iNaturalist therefore forces its users to constantly discern which organisms they consider noteworthy and which can be ignored. This process reveals the human element of choice.

To obtain estimates of birder interest for 473 bird species, we modeled variation in iNaturalist reports across species relative to the eBird baseline. To increase confidence that any findings reflect true differences in reporting behaivors, we included four critical layers of statistical rigour intended to estimate unbiased reporting rates and accurately estimate uncertainty: a spatially explicit model, a quasi-binomial error distribution, a parametric bootstrap on the overreporting index, and a phylogenetically explicit meta-analysis.

While preparing this manuscript for submission, we became aware of independent work conducted simultaneously by Callaghan et al. (12) who answer a similar question using different statistical methods and an alternative overreporting estimate.

In the wild, opportunistic birders engage more with species whose traits have been identified as charismatic in artificial contexts such as surveys (13; 16). In particular, this study reinforces the concept of the charismatic megafauna, the idea that larger species are more interesting or sympathetic to the public (32; 17; 34). The finding that more range-restricted birds are overrepresented is also consistent with the hypothesis that iNaturalist users may optimize for lengthening their “life list”, the list of unique species they have ever observed on the app(39). This “gotta catch ‘em all” commodification of nature can be at odds with the scientific pursuits that this data informs (2). The strong relationship we find between size and overreporting is also consistent with the hypothesis that iNaturalist users are drawn to birds that are easier to photograph. Logistical constraints around photography are likely not the only drivers of variation, as hinted by the fact that the American crow, a common, large, and relatively bold bird, is quite easy to photograph but is strongly underreported.

Nature photography and film-making have played a large role in conservation by bringing biodiversity to the attention of the public (5; 52; 24). As camera lenses and photos have become ubiquitous in our culture, conservation sites and museums have leveraged this fact, using Instagram and other social media platforms to understand and further engage visitors, both by producing aesthetically compelling imagery and by encouraging visitors to take and share their own photographs (59; 55; 26). iNaturalist takes advantage of commonplace camera phones to provide users with automatic identifications of their observations (49) and a platform on which to document and share their experiences. The logistical and charisma aspects of this interest bias, seen through a camera lens, are likely correlated, and we were not able to disentangle them within this framework. However, because photographs and imagery play a large role in modern communication and social media, logistical obstacles to photographing a bird may play a large role in how that bird is known or perceived by community members and therefore patterns in logistical bias may themselves be of interest. Quantifying the difference between these two community science platforms may also be of interest to scientists using the data from these sources, and future work could include leveraging these differences in modeling approaches.

It is important to note that both eBird and iNaturalist are not uniformly accessible across user demographics, so these results can only reveal the preferences of people who are already engaging with nature through these community science platforms, and who have self-selected based on a general interest in birds. Future work could investigate the “interest” filter in further sub-population analyses (e.g. children, urban areas) (6; 3) or with respect to seasonality (42).

Understanding which birds are the most charismatic can aid conservationists in creating impactful outreach materials, whether it be due to their size, color, rarity, or myriad other cultural and species-specific factors. Conservationists may tailor outreach materials, arguments, and experiences to best align with people’s preferences. This knowledge is also useful for engaging new naturalists who may not be represented in the initial analysis; by knowing which birds people tend to most enjoy seeing, an organizer can tailor a birding trip to include the most charismatic species.

## Acknowledgements

Thanks to S. Beissinger, C. Boettiger, M. Chapman, J. Clare, and W. Fithian for comments and support, and to the dedicated users of eBird and iNaturalist who provided data for this study. SS was supported by the Gordon and Betty Moore Foundation through Grant GBMF3834, and by the Alfred P. Sloan Foundation through Grant 2013-10-27 to the University of California, Berkeley. BG was supported by the National Science Foundation Graduate Research Fellowship under Grant No. 1752814. This work used the Extreme Science and Engineering Discovery Environment (XSEDE), which is supported by National Science Foundation Grant No. ACI-1548562.

## Appendix

The nine species dropped due to convergence issues in the first stage were: chestnut-sided Warbler, double-crested cormorant, Swainson’s thrush, red cross-bill, black-throated blue warbler, western kingbird, Nashville warbler, ash-throated flycatcher, and Cassin’s kingbird.

Figure 1 helps motivate the exclusion of the small number of species that had extremely negative overreporting indices. Without these values, the distribution of overreporting indices is less dramatically left-skewed.

**Figure 1.**
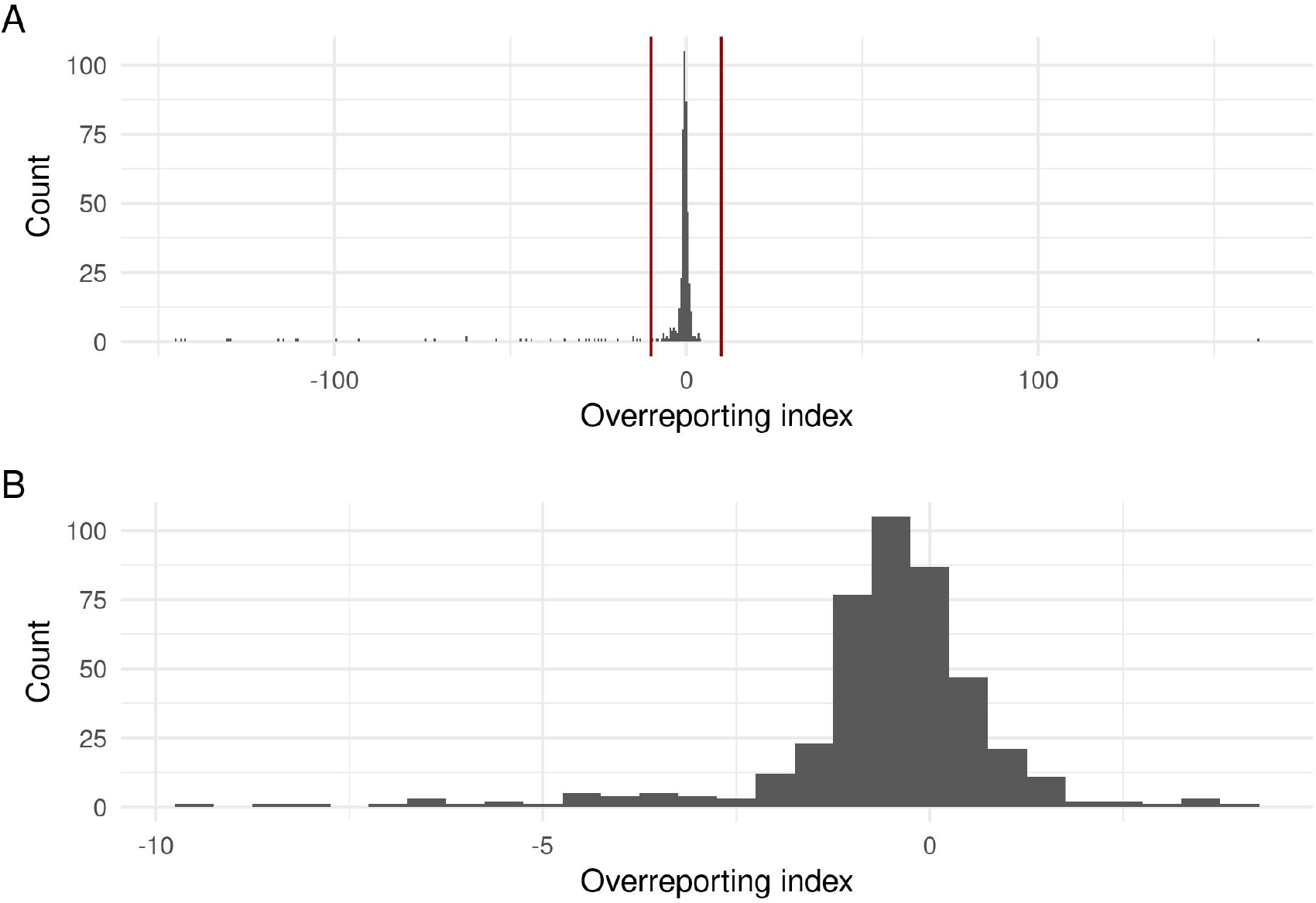
Histograms of overreporting indices before and after filtering show that the bulk of the estimates are distributed near 0, with extreme values being spread over a large range. Most, but not all, of these extremely negative values were associated with large uncertainties. Red lines indicate the cutoff (−10,10). See the Materials and methods section in the main manuscript for a discussion of how to interpret these extremely negative values.

Table 1 aims to aid the interpretation of results from the cross-species metaanalysis when low-information outliers were excluded. Mass, color, and the proxy for species range were all significant. The effects for mass and color are positive while the effect for the species range proxy is negative.

The meta-analysis results are impacted by the inclusion of the 49 low information outliers. Table 2 aims to aid the interpretation of results from the meta-analysis when low information outliers were included. The significance of color is lost and the significance of the proxy for overall prevalence is gained. The signs for mass and proxy for spatial range do not change though they get much more extreme.

**Table 2.** Meta-analysis coefficients and interpretations for species traits. All but color are statistically significant, but the effect sizes are very extreme on this scale, alluding to instability.

| Covariate | Effect Size | Interpretation |
| --- | --- | --- |
| Scaled log mass | 4.81 (0.49, 9.14) | An increase by one standard deviation of bird size on the log scale is associated with an expected nearly 23-fold increase in overreporting. |
| Scaled color | 0.23 (-1.47, 1.92) | An increase by one standard deviation of bird color contrast is associated with an expected 26% decrease in the odds of overreporting. |
| Scaled log number of hexes where spotted | -4.35 (-6.50, -2.19) | An increase by one standard deviation of number of hexes on the log scale is associated with an expected nearly 78-fold decrease in the odds of overreporting. |
| Scaled log proportion of eBird checklists where found | 5.96 (3.61, 8.31) | An increase by one standard deviation of proportion of eBird checklists where a species is found, on the log scale, is associated with a nearly 388-fold increase in overreporting. |

The 49 low-information outliers were: Abert’s towhee, alder flycatcher, Arctic tern, Bachman’s sparrow, bank swallow, Bicknell’s thrush, black rosy-finch, black swift, black-chinned sparrow, boreal owl, Cassin’s vireo, Connecticut warbler, Cordilleran flycatcher, dusky flycatcher, Eastern whip-poor-will, elf owl, flammulated owl, gray flycatcher, greater sage grouse, Hammond’s flycatcher, Henslow’s sparrow, hepatic tanager, hoary redpoll, Hudsonian godwit, Kentucky warbler, king eider, king rail, Lucy’s warbler, mountain plover, mountain quail, Nelson’s sparrow, parasitic jaeger, Philadelphia vireo, pinyon jay, red knot, red phalarope, red-cockaded woodpecker, sage thrasher, Smith’s longspur, Stejneger’s scoter, stilt sandpiper, Swainson’s warbler, Vaux’s swift, Virginia’s warbler, whiteheaded woodpecker, Williamson’s sapsucker, yellow rail, yellow-bellied flycatcher, and yellow-billed loon.

These low-information outliers are generally extreme overreporting indices. These primarily arise from cases of few or no iNaturalist observations in many of the hexes where the bird was reported in eBird. This could be partially explained by the fact that iNaturalist has far fewer total bird observations such that many rare or underreported species will never appear in the dataset in certain hexes, and therefore a rate difference cannot be reliably estimated in spite of the fact that a parametric bootstrap of median differences consistently predicts an extreme difference.

## Notes

### Competing Interest Statement

The authors have declared no competing interest.

